# Compartment and cell type-specific hypoxia responses in the developing *Drosophila* brain

**DOI:** 10.1101/2019.12.12.874404

**Authors:** Martin Baccino-Calace, Daniel Prieto, Rafael Cantera, Boris Egger

**Affiliations:** Developmental Neurobiology, Instituto de Investigaciones Biológicas Clemente Estable, Montevideo 11600, Uruguay; Institute of Molecular Life Sciences, University of Zürich, Zurich CH-8057, Switzerland; Zoology Department, Stockholm University, Stockholm 106 91, Sweden; Department of Biology, University of Fribourg, Fribourg CH-1700, Switzerland

## Abstract

Environmental factors such as the availability of oxygen are instructive cues to regulate stem cell maintenance and differentiation. We used a genetically encoded biosensor to monitor the hypoxic state of neural cells in the larval brain of *Drosophila*. The biosensor reveals brain compartment and cell type specific levels of hypoxia. The values correlate with differential tracheolation that is observed throughout development between the central brain and the optic lobe. Neural stem cells in both compartments show the strongest hypoxia response while intermediate progenitors, neurons and glial cells reveal weaker responses. We demonstrate that the distance between a cell and the next closest tracheole is a good predictor of the hypoxic state for that cell. Our model concludes that oxygen availability is the major factor controlling the hypoxia response in the developing *Drosophila* brain but cell intrinsic and cell-type specific factors contribute to modulate the response in an unexpected manner.

## INTRODUCTION

In stem cell niches the supply of oxygen and nutrients is tightly controlled (Schofield, 1978; Morrison and Spradling, 2008) and the stem cell microenvironment ensures a balanced response of stem cells to the needs of the organism (Li and Xie, 2005; Scadden, 2006; Jones and Wagers, 2008). It was reported that embryonic, hematopoietic, neural and cancer stem cells reside in hypoxic niches (Ezashi et al., 2005; Simon and Keith, 2008; Mohyeldin et al., 2010; Simsek et al., 2010; Takubo et al., 2010; Lee and Simon, 2012) and that hypoxia favours survival, maintenance and proliferation of stem cells *in vitro* and *in vivo* (Csete, 2005; Carnero and Lleonart, 2016).

The first studies reporting a functional relationship between neural precursor cells and hypoxia was performed in the carotid body, where glomus cells showed increased survival and proliferation in response to hypoxia (Bee et al., 1986; Nurse and Vollmer, 1997; Pardal et al., 2007). Hypoxia increases multipotency, proliferation and selective survival of neural stem cells. On the contrary, exposing stem cells to atmospheric oxygen causes differentiation and cell death (Studer et al., 2000; Storch et al., 2001; Gustafsson et al., 2005; Pardal et al., 2007; Panchision, 2009). More recently, Lange and colleagues demonstrated that the relief of tissue hypoxia by ingrowing blood vessels is an instructive signal for neural stem cell differentiation in the developing cerebral cortex (Lange et al., 2016).

In the *Drosophila* embryo a fully functional nervous system is built within a few hours of embryonic development, which serves the freshly hatched larva among other behaviours to navigate and feed. During larval growth, a second wave of neurogenesis is initiated to produce the neurons for the adult central brain and the ganglia of the optic lobes (Truman and Bate, 1988; Hofbauer and Campos-Ortega, 1990; Green et al., 1993; Meinertzhagen and Hanson, 1993; Hartenstein et al., 2008). As the stem cells of the optic lobes proliferate, their progenies await in an arrested state of differentiation for several days before they become fully differentiated and form synaptic connections in mid-pupal life (Melnattur and Lee, 2011; Chen et al., 2014). Hence, in the larval optic lobe proliferating progenitor cells co-exist during several days with postmitotic cells that remain in a state of arrested differentiation.

As the nervous system develops in the embryo, tracheal cells invade the brain along the dorsal midline and build the network of respiratory tubes, called tracheoles that oxygenate the brain during larval life. In *Drosophila* these air tubes have a stereotyped branching pattern, making it possible to draw a detailed map of the larger tracheoles reaching each brain region (Pereanu et al., 2007). The present study was prompted by the observation that in the developing larval brain tracheoles are not distributed as homogeneously and densely as in muscle, ovary, intestine and other tissues with high metabolism (Bownes, 1982; Li et al., 2013; Peterson and Krasnow, 2015; Misra et al., 2017). In the larval brain the tracheal network is largely segregated into two main compartments. A central region, where the functional neuronal circuits are located, is densely tracheolated. However, to each side of the central brain there is a large compartment containing very few tracheoles (Misra et al., 2017). These lateral regions correspond to the proliferative anlagen of the optic lobes (White and Kankel, 1978; Hofbauer and Campos-Ortega, 1990).

Our main hypotheses are that the sparse tracheolation of the optic lobes is an essential aspect of normal brain development because it results in a state of constitutive hypoxia, relative to the central brain, which in turn will promote proliferation and inhibit differentiation of the newly formed neurons. Quantitative data obtained with a hypoxia biosensor supports the notion that the optic lobe is less oxygenated than the central brain (Misra et al., 2017).

Here we mapped the hypoxic states of different brain regions throughout larval development and found that the proliferative anlagen of the optic lobes show elevated hypoxia levels as compared to the densely tracheolated and synaptically active central brain. The high spatial resolution of the biosensor made it possible to detect consistent differences in the hypoxia values assigned to cells located very close to each other, and evidence is presented for cell type-specific hypoxia responses. We analysed the relationship between tracheolation and the hypoxia response revealed by the biosensor. Interestingly, we find that the minimum distance between a cell and the next tracheole is a good predictor of the hypoxic state of that cell. Finally, we provide evidence that neural progenitor cells respond to altered ambient oxygen levels in a cell-type specific manner. We conclude that this knowledge opens the opportunity to use *Drosophila* for the study of how hypoxia regulates stem cell proliferation and neuronal differentiation. Knowing what factors control the co-existence of proliferating tissue within a differentiated organ such as the brain is also of interest for studying tumour formation and maintenance.

## RESULTS

### Differential tracheolation persists throughout larval development

In the developing larva of *Drosophila*, the central brain is much more densely tracheolated than the optic lobe. Using a novel hypoxia biosensor, we found that towards the end of larval life this asymmetry in the density of tracheoles correlates with asymmetry in hypoxia levels, with the optic lobe being less oxygenated than the central brain (Pereanu et al., 2007; Misra et al., 2017). Here, we used confocal laser microscopy and transmission electron microscopy to further define this morphological and functional dichotomy and investigated whether this condition prevails throughout larval development. In brains immunolabelled for the synaptic marker Bruchpilot (Brp) (Kittel et al., 2006; Wagh et al., 2006) the staining co-localizes with regions of the brain that are densely tracheolated (Figure 1A, B) and that correspond almost entirely to the synaptic centres (i.e. neuropils, see for example (Iyengar et al., 2006)). In contrast, very little synaptic staining was found within the optic lobes (Figure 1B). Hence, there is a close topographic correlation between a dense tracheolation of the synaptic neuropil in the central brain and a sparse tracheolation in the optic lobes, where there are no synapses but instead undifferentiated progenitor cells (Figure 1C). A close examination of the border between these two brain regions, using transmission electron microscopy disclosed the existence of a sharp interphase between two types of cell bodies (inset in Figure 1C and 1D). On the side of the central brain we found glia, tracheoles and neuronal cell bodies extending thick neurites into the neuropil where they formed synapses (not shown). These cell bodies had a relatively large cytoplasm containing abundant mitochondria, endoplasmic reticulum, ribosomes and other organelles, as expected for differentiated neurons (Figure 1E). On the opposite side (Figure 1F), within the medial region of the optic lobes, we found large numbers of smaller cells, in which the nucleus was surrounded by a thin ring of cytoplasm, with fewer organelles and with the typical columnar arrangement of the yet not fully differentiated neuronal progeny generated in this proliferative region (Meinertzhagen and Hanson, 1993; Fischbach and Hiesinger, 2008; Hasegawa et al., 2011; Melnattur and Lee, 2011).

**Figure 1.**
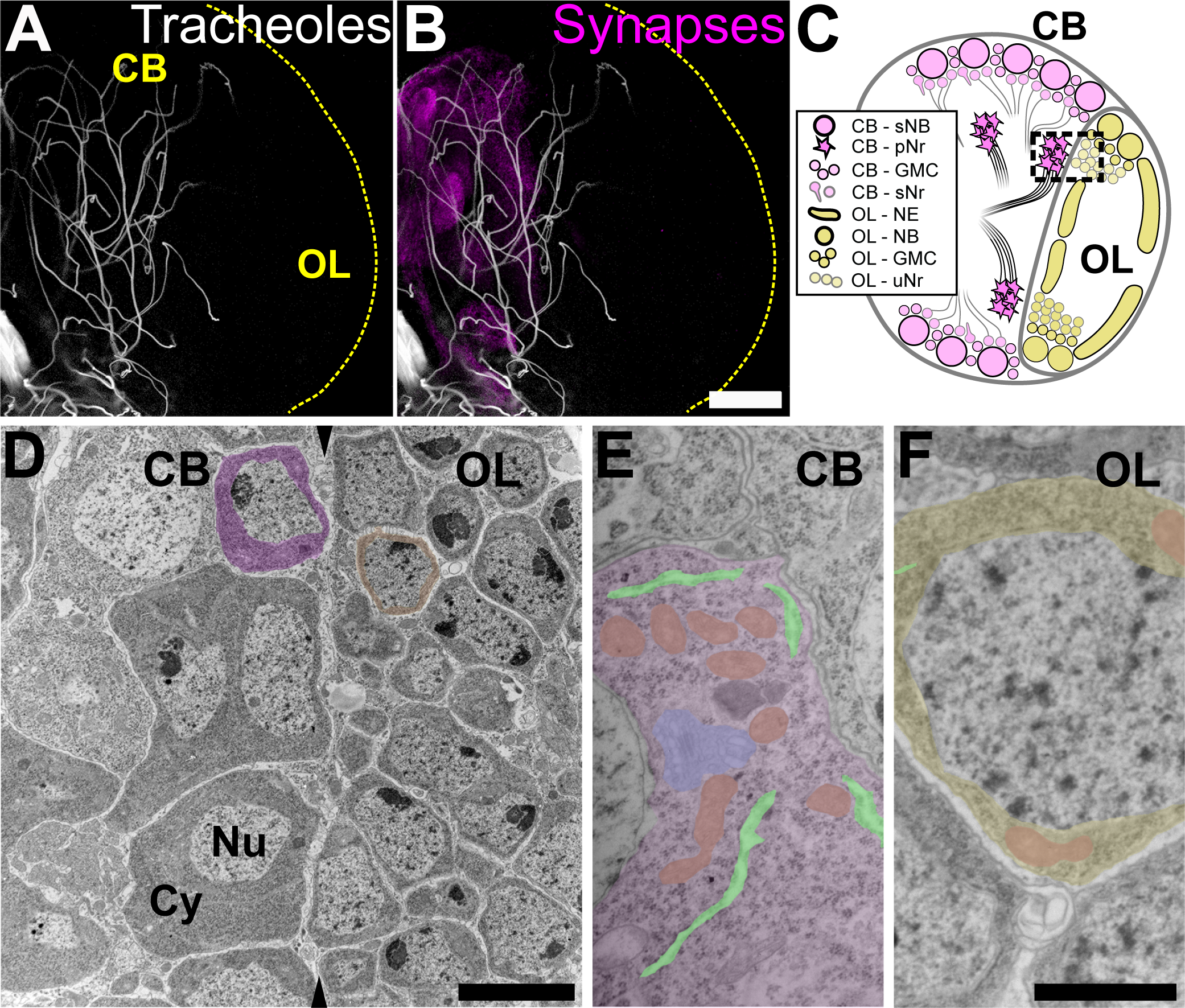
Tracheoles and undifferentiated cells are spatially segregated within the larval brain. Tracheoles (here imaged by means of their autofluorescence) are abundant within the central brain (A, left part of the hemisphere) as well as Bruchpilot-positive synapses (B), but sparse in the optic lobe (right part of the hemisphere, see also C) (n=6). The larval optic lobe is enriched in immature cells whilst most terminally-dfferentiated neurons are found in the central brain. At some planes a boundary can be observed between neurons and progenitors (C, dashed square). Transmission electron microscopy of a region similar to that squared in C reveals the boundary (arrowheads in D) between central brain differentiated neurons with a low nucleus/cytoplasm ratio (left in D, pseudo-coloured in magenta) and undifferentiated progenitors from the optic lobe, with much less cytoplasm (right in D, pseudo-coloured in yellow) (n=5). Neurons show big cell bodies enriched in organelles like mitochondria (E, pseudo-coloured in orange), rough endoplasmic reticulum cisternae (pseudo-coloured in light green), and Golgi cisternae (pseudo-coloured in cyan) and other organelles. Progenitor cells from the optic lobe (F) are less differentiated than primary central brain neurons and show a thin ring of cytoplasm around the nucleus. CB: central brain; OL: optic lobe; Nu: nucleus; Cy: cytoplasm; sNB: secondary neuroblasts; pNr: primary neurons; GMC: ganglion mother cells; sNr: secondary neurons; NE: neuroepithelium; NB: neuroblasts; uNr: undifferentiated neurons. Scale bar B-C: 50 µm; D: 5 µm; E-F: 500 nm.

We previously reported that at a late stage of larval development, 96 hrs after larval hatching (ALH), the sparsely tracheolated optic lobe has lower hypoxia values than the densely tracheolated central brain (Misra et al., 2017). To investigate if this condition is specific for the end of larval life or prevails during a longer developmental interval and is thus of potential relevance for brain development we extended our analysis to seven time points of larval development at 12 hr intervals, from 24 to 96 hrs ALH. We found that the segregation of tracheoles exists already by 24 hrs ALH (Figure 2A) and persists throughout larval life (Figure 2A-F). We confirmed that the optic lobe grows in size during this time (Figure 2G) (mean values in µm^3^, 24hrs: 28206.5, 36hrs: 105438.5, 48hrs: 85418.4, 60hrs: 381289.3, 72hrs: 1696572.2, 84hrs: 1909378.7, 96hrs: 2271840.1, n=4, 6, 6, 6, 6, 5,6, respectively). We observed that also the tracheoles grow considerably in overall length (Fig. 2H; values in µm, 24hrs: 76.2., 36hrs: 144.7, 48hrs: 266.8, 60hrs: 503.0, 72hrs: 625.3, 84hrs: 511.6, 96hrs: 684.3) although not enough to compensate for optic lobe growth from 48h ALH onwards, and thus the proportion of optic lobe tissue devoid of tracheoles appears to increase with age (Figure 2I).

**Figure 2.**
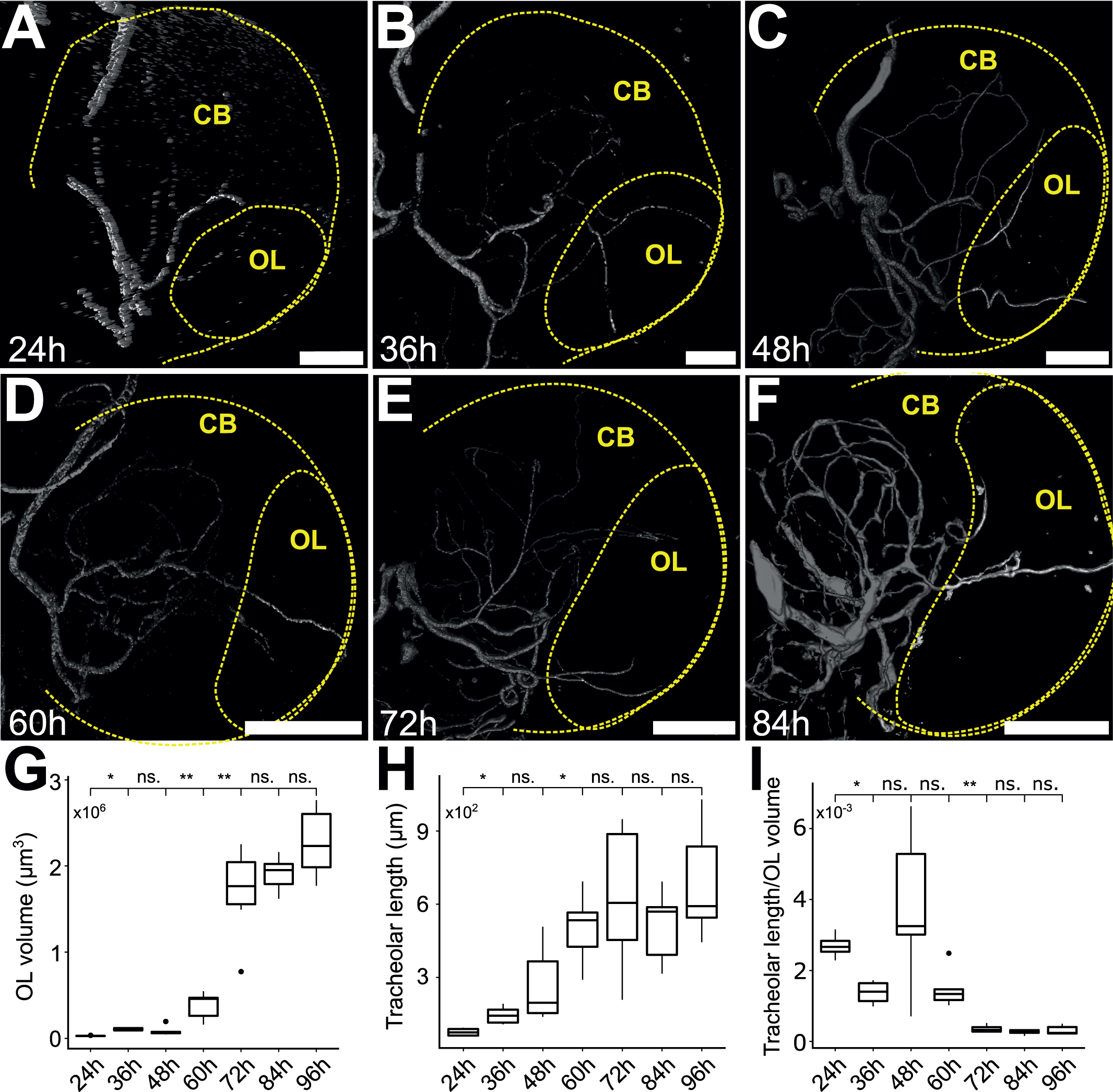
The volume and the length of tracheoles of the optic lobe increase during larval development. Tracheal trees reconstructed from calcofluor white staining of 24-84h ALH larval brains at 12h intervals (A-F). The hemisphere and optic lobe are outlined with dashed lines. Optic lobe volume measurement from 24-96h ALH larval brains at 12h intervals show continuous growth (G), as well as total tracheolar length within the optic lobe (H). Analysis of the ratio of tracheal length and optic lobe volume show that growth in optic lobe volume is not compensated by normal increase of tracheal length (I). Scale bars are 20 µm for panels A-C, 50 µm for panels D-F. * p<0.05, ** p<0.001, Mann-Whitney Wilcoxon test. ns.: non-significant; ALH: after larval hatching. Sample sizes for time points from 12hrs to 96hrs ALH n = 4, 6, 6, 6, 6, 5, 6.

### Oxygen availability triggers differential hypoxia response between central brain and optic lobe

Since our original hypothesis stated that the sparse tracheolation of the optic lobes will result in a condition of chronic hypoxia relative to the central brain, we used a HIF-1α/Sima based hypoxia sensor (Misra et al., 2017) to monitor hypoxia in these two brain compartments throughout larval life. The results were consistent with this prediction because mean biosensor ratiometric values were significantly lower for the optic lobe (stronger hypoxia response) as compared to the central brain at 36, 60 and 84 hrs ALH (Figure 3; mean ratiometric values at 36 hrs: 0.90 for central brain, 1.23 for optic lobe, n=6; at 60 hrs ALH: 0.89 for central brain, 1.22 for optic lobe, n=6; at 84 hrs ALH, 0.90 for central brain, 1.31 for optic lobe, n=8). Our results presented here so far strongly suggest that the dense tracheolation of the central brain results in higher oxygenation of this compartment in comparison with the sparsely tracheolated optic lobe.

**Figure 3.**
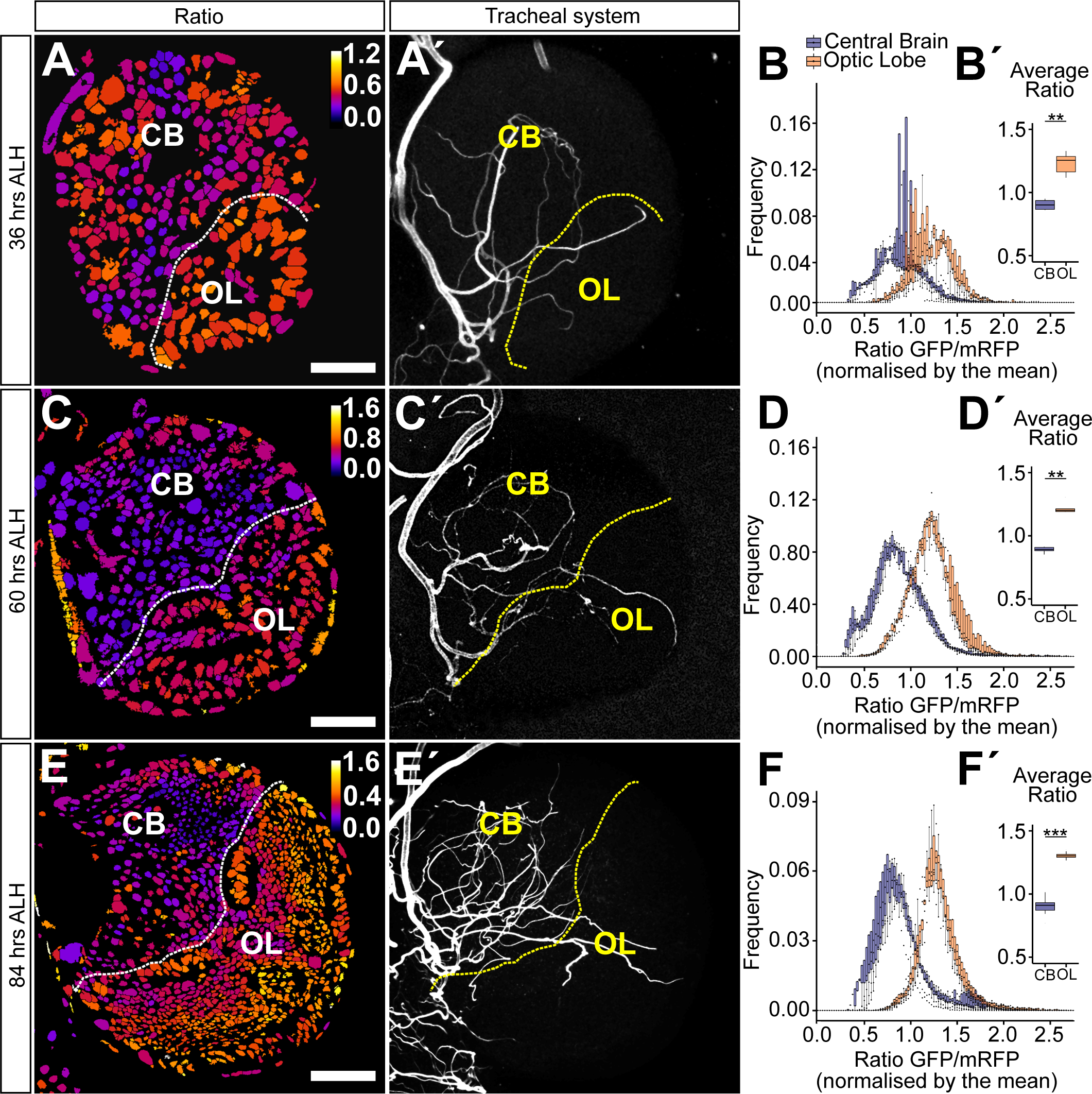
Hypoxia response is differentially regulated between central brain and optic lobe throughout larval development. (A, C, E) ratiometric images of single frontal confocal section of a brain hemisphere of a larva expressing green (ubi-GFP-ODD) and red (ubi-mRFP-nls) fluorescent proteins of the biosensor. The colour code (upper right) indicates average GFP-ODD/mRFP-nls ratios for each nucleus, which were segmented based on the mRFP-nls signal. (A’, C’, E’) maximum intensity projection of the brain tracheal system (white). The dotted line denotes the separation between central brain and optic lobe. (B, D, F) histograms representing the frequency distribution of GFP-ODD/mRFP-nls ratios (normalised to whole brain average) for the central brain and the optic lobe, showing a clear difference in the distribution of values between the two compartments. (B’, D’, F’) box plots showing the non-normalised mean GFP-ODD/mRFP-nls ratios. Box plots in (B’, D’, F’) show maximum and minimum observation, upper and lower quartile, and median. Scale bars are: 15 µm (A, A’), 20 µm (C, C’), 40 µm (E, E’). * p<0.05, ** p<0.01, *** p<0.001 student t-test or Mann-Whitney Wilcoxon test. Sample size n= 6, 6, 8.

### The distance between a cell and its nearest tracheole can predict its cellular hypoxia response

To further investigate the relationship between tracheolation and the distribution of oxygen in the brain we decided to focus on the lateral optic lobe tracheole (OLTl) described by Pereanu and collaborators (Pereanu et al., 2007), which enters the optic lobe through the inner proliferation center (IPC). The spatial separation of the OLTI from other tracheoles provides a good opportunity to analyse how oxygen availability and hypoxic response values change within a group of similar cells, which only differ in their distance to a neighbouring tracheole. Our measurements indicate that as the minimal distance between cells and OLTl increases, hypoxia values increase. It suggests that oxygen availability decreases (Figure 4). The data (1/Ratio as a function of distance) best fits a decaying exponential function of the shape:

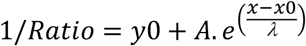

**Figure 4.**
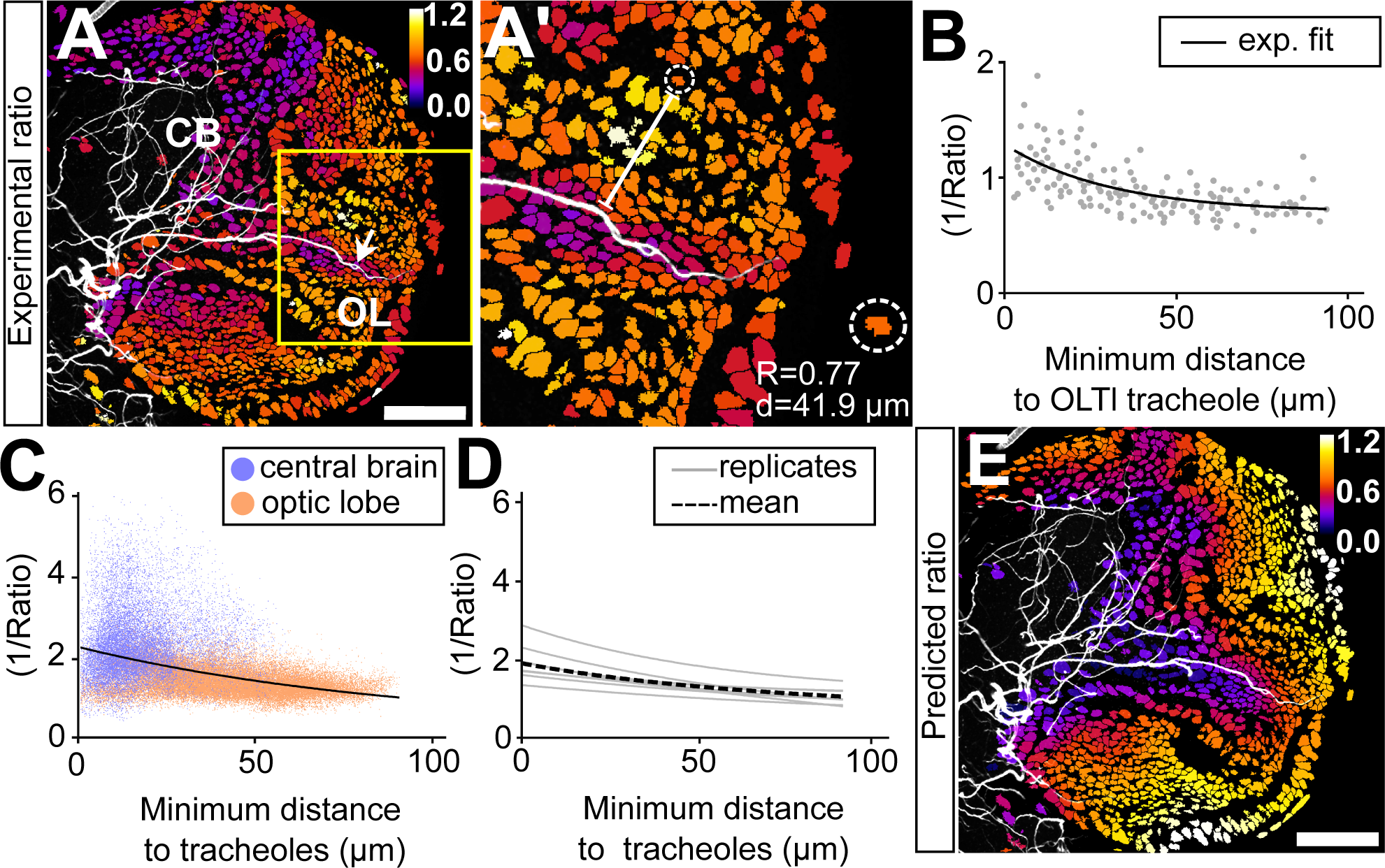
Hypoxia response correlates with distance between a cell and closest next tracheole. (A) ratiometric image of an larval hemisphere 84 hrs ALH superimposed with a maximum projection of the tracheal system (white). (A’) magnified view of yellow square in (A) which points to nuclei from the optic lobe volume surrounding the optic lobe lateral tracheole (OLTl, white arrow in (A)). Inset shows a single nucleus (inside dotted circle) for which the corresponding ratio value and distance to tracheole are shown (bottom right). (B) 1/Ratio plotted against distance to OLTl tracheole. Black line shows exponential fit to the data (n=6). (C) 1/Ratio plotted against minimum distance to tracheole for every nucleus in the brain (purple: central brain, orange: optic lobe; n=6). Black line is an exponential fit to the data. (D) exponential fits (red lines) for values of 8 different brains; black dotted line shows average fit for these brains. (D) the function resulting of the exponential fit to the data was used to calculate a predicted ratio value if such value was solely based on the distance of a given nuclei to the closest tracheole. Scale bar: 40 µm. These values where depicted with a colour code in (E).

(Figure 4B). The study of the OLTl tracheole provided key insights into the way oxygen diffuses from a given tracheole and prompted us to apply the same analysis in a generalized way to the entire brain hemisphere. We produced a map that contains the coordinates of every cell in the brain and measured the minimum distance between each cell and neighbouring tracheoles in the entire brain. 1/Ratio values plotted as a function of minimum distance to trachea show an inverse relationship that is best fitted by a decaying exponential function (Figure 4C and 4D). The result demonstrates that the hypoxia response of individual cells correlates with their position in the brain in relation to the tracheal system. Using the best fit exponential function, we predicted ratiometric values according to cell-to-tracheole distance and depicted them with a colour scale in a heat map (Figure 4E). The image strongly resembles the distribution of measured ratiometric values (compare Figure 4E to 4A), which indicates that the distance to tracheoles can reliably predict the hypoxic, or conversely, the oxygenation state of cells.

However, the biosensor might not distinguish between the contribution of oxygen availability and other factors that affect HIF-1α/Sima and ODD-GFP degradation. In order to demonstrate that oxygen is the main determinant of the differences of ratiometric values, we subjected larvae to altered ambient oxygen conditions (Figure 5). We exposed larvae to increased atmospheric oxygen levels (hyperoxia) by raising them in 60% oxygen from 24 hrs to 84 hrs ALH (Figure 5A-C). The results were consistent with an increase in oxygen dependent degradation of GFP-ODD in the optic lobe, relative to the central brain, indicating enhanced oxygen availability. The frequency distribution of ratiometric values showed that optic lobe values were shifted towards central brain values as compared with brains from larvae kept in normoxia (compare Figure 5B and 3F). Hence, the difference in the average hypoxia response between the central brain and the optic lobe decreased significantly upon ambient hyperoxia (Figure 5B’; mean ratiometric values for central brain in normoxia: 0.89, optic lobe in normoxia: 1.32; central brain in hyperoxia: 0.55, optic lobe in hyperoxia: 0.61, n=7). Under these conditions, cells of the central brain and optic lobe seem to experience similar oxygen levels despite the persistence of unequal tracheolation in these two compartments (Figure 5A, A’). The decay kinetics of oxygen levels (1/Ratio) in relation to distance to tracheoles was reduced, which indicates oxygen saturation of the brain and the loss of the difference between central brain and optic lobe (Figure 5C; normoxia: black curve, λ: 242 µm, hyperoxia; blue line, λ: 653 µm).

**Figure 5.**
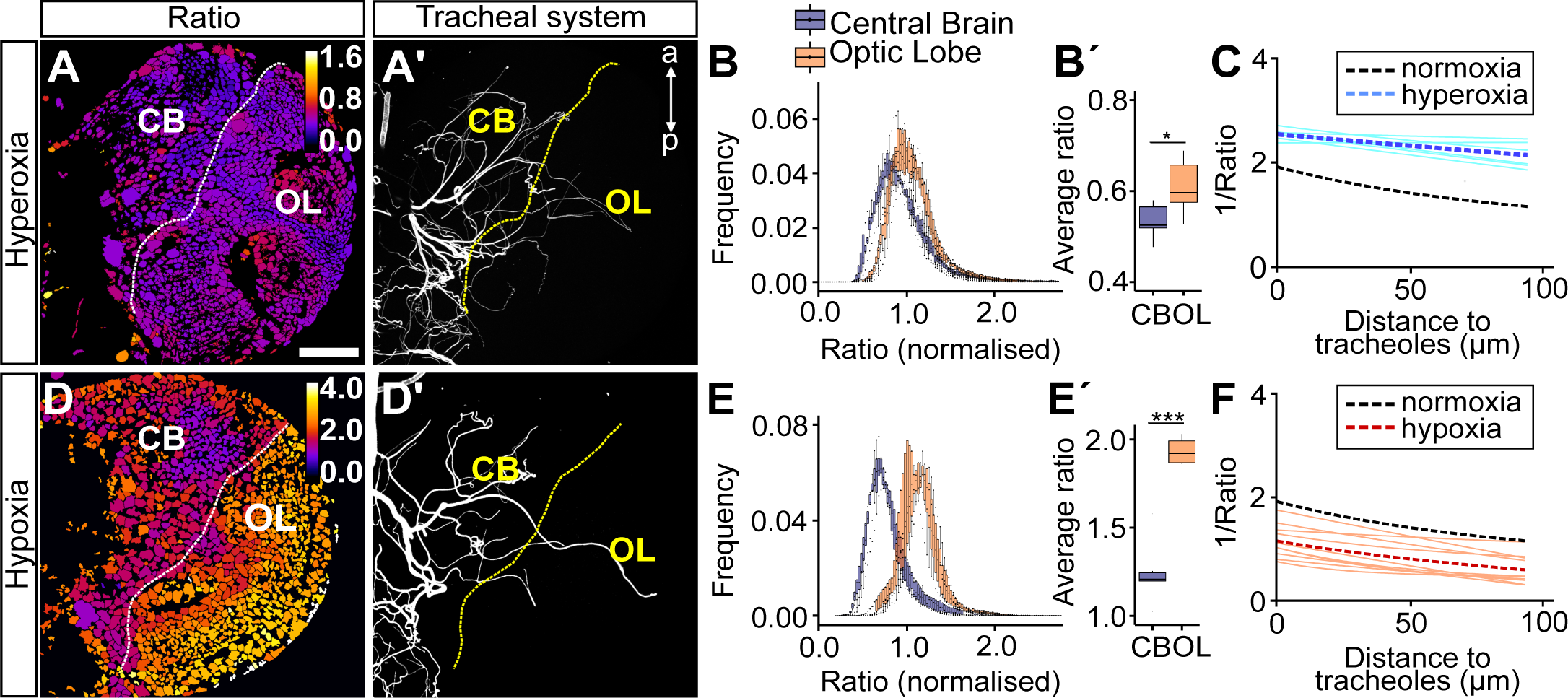
GFP-ODD degradation is driven by oxygen availability. Ratiometric images for larva exposed to hyperoxia (A, n=7) and hyperoxia (D, n=7) and their corresponding maximum projection of their tracheal system (A’, D’). Larva reared in ambient hyperoxia show a left-shift (higher oxygen) in the distribution of optic lobe ratio values (B). Brains of larva exposed to hyperoxia show lower ratio values both for central brain and optic lobe (B’). Larva reared in hypoxia show two distinct populations of values for central brain and optic lobe (E) as observed in normoxia (see Figure 3E). (E’) non-normalized mean ratio values are increased (lower oxygen) as compared to normoxia. (C, F) 1/Ratio (oxygenation) plotted against minimum distance to trachea. Sky blue lines (C) and orange lines (F) show exponential fits for different brains from larva reared in hyperoxia and hypoxia, respectively. Dotted lines show average of all fits for normoxia (black), hyperoxia (dark blue) and hypoxia (red). Scale bars are 40 µm. * p<0.05, ** p<0.01, *** p<0.001 student t-test or Mann-Whitney Wilcoxon test.

Next, we exposed larvae to ambient hypoxia by raising them in 5% oxygen from 60 hrs to 84 hrs ALH in order to investigate whether the sensor is able to show decreased oxygen availability in the brain. We observed an increase in ratiometric values in both central brain and optic lobe compartments (Figure 5D, E’). The difference in the average hypoxia response between central brain and optic lobe increased significantly in larvae kept for 24 hrs in ambient hypoxia (Figure 5E’; mean ratiometric values for central brain in normoxia: 0.89, optic lobe in normoxia: 1.3, central brain in hypoxia: 1.25, optic lobe in hypoxia: 1.86, n=7). The relationship between oxygenation (1/Ratio) and distance to tracheoles remained unchanged compared to normoxia (Figure 5F; normoxia: black curve, λ: 242 µm hypoxia: red line, λ: 414 µm).

Overall these results support the notion that the differences in ratiometric values observed in our experiments are predominantly due to oxygen tensions and not regulated by other factors influencing HIF-1α/Sima degradation. However, closely studying the map of hypoxia-response predicted values (Figure 4E) we noticed that for certain cell types these predicted values deviated from the measured values (Figure 4A). We therefore analysed the hypoxia response in more detail by analysing the biosensor in a cell type specific manner

### Biosensor reveals cell-type specific hypoxia states in central brain and optic lobe

The results of our brain-compartment analysis prompted us to investigate the hypoxia response in different cell types found in the central brain and in the optic lobe (Figure 6). For this we combined the ratiometric analysis with immunofluorescence labelling for cell type specific nuclear marker proteins. We marked neuroblasts with an antibody against Deadpan (Dpn), ganglion mother cells (GMCs) with an antibody against Prospero (Pros), neurons with an antibody against Embryonic lethal abnormal visual system (Elav) and glial cells with an antibody against Reversed polarity (Repo) (Figure 6A-D). Neuroepithelial cells were segmented based on a staining with an antibody against Disc large (Dlg) using TrakEM2 in Fiji (Figure 6E). Image stacks obtained from the immunostainings with these markers served to produce cell type-specific segmentation masks for the ratiometric analysis (Figure 6A’-E’).

**Figure 6.**
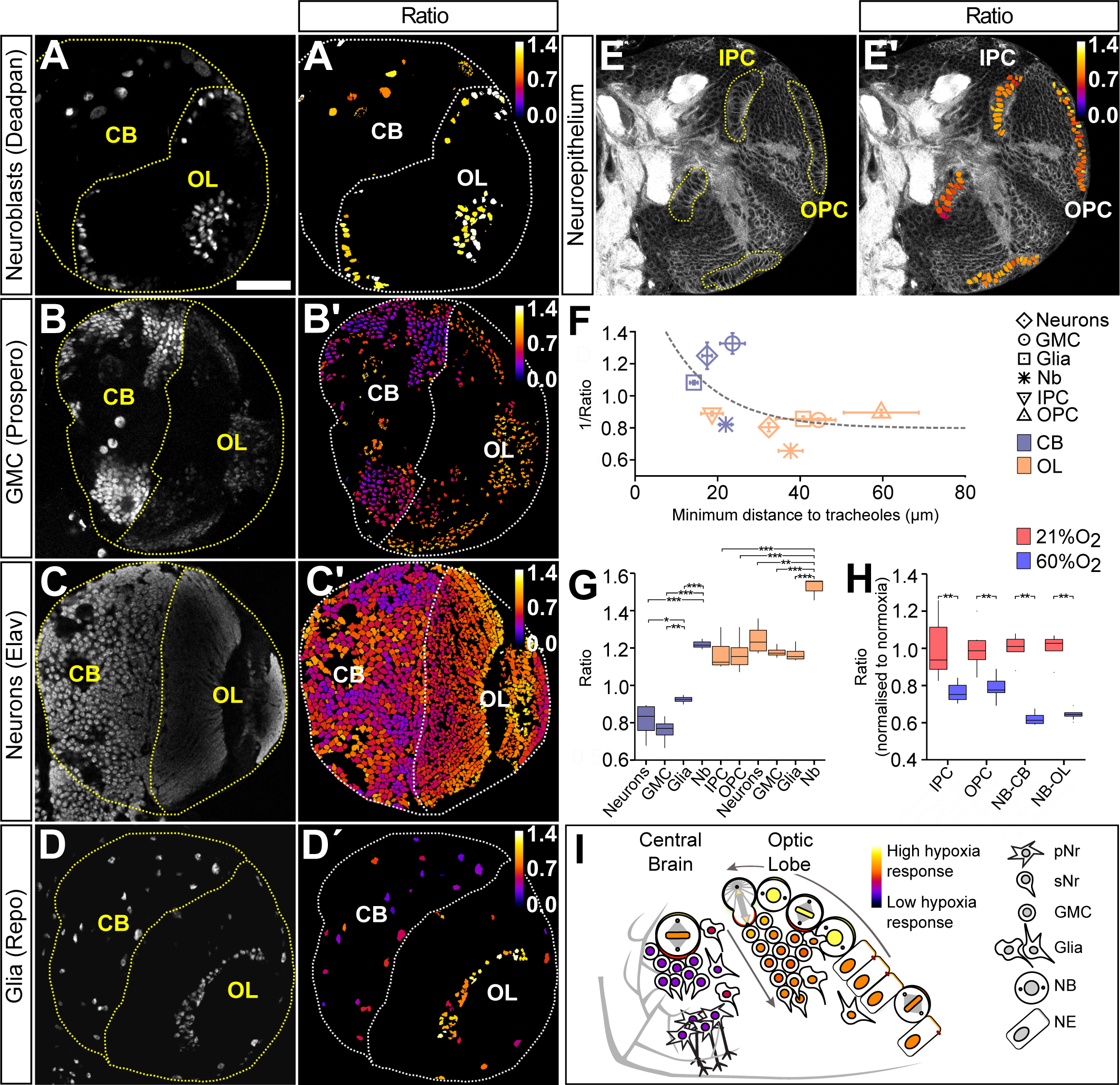
The biosensor reveals cell-type specific hypoxia states in central brain and optic lobe. (A to E) immunostainings for cell-type specific markers (gray): (A) anti-Deadpan (Neuroblasts, Nb), (B) anti-Prospero (Ganglion mother cells, GMC), (C) anti-Elav (neurons), (D) anti-Repo (glial cells) and (E) anti-Disc large (used to segment neuroepithelial cells). The morphology of neuroepithelial cells (IPC and OPC) allowed us to manually segment these cell populations using TrakEM2 based on the Disc large signal. The images from (A to D) were utilized as segmentation signals to create a mask for cell-type specific ratiometric analysis (A’ to E’). (F) mean 1/Ratio (oxygenation) for central brain and optic lobe for each cell type is represented as a function of distance to trachea. The dotted line shows an exponential fit to the data points. (G) Boxplot compares ratiometric values for all cell types both in central brain and optic lobe (n=4). (H) Exposure to hyperoxia has a stronger effect in neuroblasts than in neuroepithelial cells of the IPC and OPC (n=6). (I) illustrates cell type specific hypoxia response in the central brain and optic lobe based on biosensor data. Scale bar 40 µm. Error bars in (F) show S.E.M. * p<0.05, ** p<0.01, *** p<0.001 student t-test or Mann-Whitney Wilcoxon test.

In general, and as expected from our brain-compartment analysis, each cell type in the central brain showed a lower hypoxic response as compared to the same cell type in the optic lobe (Figure 6F). Interestingly, however, we observed cell-type specific differences in the hypoxia response between cell types, regardless of the localisation of the cells in the brain. Neuroblasts (Dpn-positive cells) in the optic lobe showed the strongest hypoxia response, while ganglion mother cells (Pros-positive cells) in the central brain showed the weakest hypoxia response of all analysed cell types in the brain (mean ratio values for optic lobe neuroblasts: 1.51 and for central brain ganglion mother cells 0.75) (Figure 6A, B and G). In both the central brain and the optic lobes, Dpn-positive neuroblasts were on average more hypoxic than any other cell type in the corresponding brain compartment. Neurons (Elav-positive cells) and glial cells (Repo-positive cells) showed intermediate levels of hypoxia response in the corresponding brain compartments (Figure 6C, D; mean ratio values for central brain neurons: 0.81, optic lobe neurons: 1.24 and central brain glial cells 0.92, optic lobe glial cells: 1.16, n=4).

We next measured the average minimum distance between cells of a defined cell type and neighbouring tracheole (Figure 6F). The average 1/Ratio values (oxygenation) for most cell types in the central brain and the optic lobe followed a decaying exponential function as described above. Of interest is that central brain and optic lobe neuroblasts showed a stronger and central brain ganglion mother cells a much weaker hypoxia response as compared to the prediction. Most unexpected based on the predicted map was the observation that neuroepithelial cells of the outer and inner proliferation centre (OPC and IPC) revealed virtually the same level of hypoxia response (Figure 6E, mean ratio values for OPC: 1.16 and IPC: 1.16, n=4). This is remarkable because IPC neuroepithelial cells are located much closer to the densely tracheolated central brain compartment than OPC neuroepithelial cells.

### Neuroepithelial cells are resilient to differential oxygen levels

In order to investigate the idea that certain cell types are less efficient than others in sensing oxygen levels via the canonical hypoxia pathway we compared the hypoxia response in neuroepithelial cells and neuroblasts under ambient hyperoxia (Figure 6H). Interestingly, while neuroblasts in the central brain and in the optic lobe were able to greatly adapt their hypoxia response to different oxygen levels, neuroepithelial cells of the IPC and OPC changed their response to a much lower degree. This indicates that neuroepithelial cells might be less susceptible to changes in oxygen levels.

This finding prompted us to investigate a dataset with genome-wide information on larval brain gene expression (Southall et al., 2013). In this study, cell-type specific targeted DamID methods were used to compare Polymerase II occupancy between neuroepithelial cells and neuroblasts in the third instar larval brain at age 96 hrs ALH. In both progenitor cell types, glycolytic genes were significantly enriched, indicative of a lower oxygen availability to use oxygen for oxidative respiration for cellular energy production. Moreover, enrichment in hypoxia pathway genes was found in the neuroblast specific gene catalogue but not in neuroepithelial cell specific gene catalogue (Table 1). The results support the notion that in neuroepithelial cells the canonical hypoxia pathway might be less relevant in order to respond to differential oxygen levels.

**Table 1.** Functional Enrichment Analysis.

| *Biological process* | *Cell type* | *Enriched* | *p-value* |
| --- | --- | --- | --- |
| Glycolysis | NE | YES | 6.9 x10<sup>-3</sup> |
|  | NB | YES | 3.9 x10<sup>-2</sup> |
| Hypoxic response | NE | NO | >0.05 |
|  | NB | YES | 9.3 x10<sup>-4</sup> |
| NE: Neuroepithelial cells.<br>NB: Neuroblasts. |  |  |  |

## DISCUSSION

Ambient oxygen has crucial roles for the development and the growth of organisms (Harrison and Haddad, 2011). Two recent studies demonstrate how *Drosophila* larvae can adapt to low atmospheric oxygen or hypoxia by reducing their growth rate through systemic mechanisms (Lee et al., 2019; Texada et al., 2019). Less understood are the direct effects of differential oxygen supply within a tissue on cellular maintenance, proliferation and differentiation. Our knowledge is sparse on how different cell types within an organ or a tissue respond to various levels of oxygen provided by the vascular or the tracheal system, in mammals and insects, respectively.

Our study is based on the observation of significant differences in tracheolation between brain compartments in the developing *Drosophila* larvae. While the central brain is densely tracheolated, the optic lobes are only very sparsely tracheolated throughout larval life. This dichotomy in tracheolation correlates with cellular differentiation within the two compartments and indicates that keeping relatively low levels of oxygen might be important for optic lobe progenitor proliferation and maintenance. Indeed, hypoxia has an important function in the neural stem cell niche and for normal neural development. A tight spatial and temporal correlation between the degree of vascularization and the segregation of proliferative and differentiating zones of the brain were found in the developing mammalian cerebral cortex. Here, the proliferative zone is poorly vascularised and stem cells rely on glycolytic metabolism for energy production. With the arrival of the first blood vessels oxygen levels are elevated, which leads to HIF1-α degradation and neuronal differentiation. Blocking angiogenesis in the proliferative zones maintains the proliferative state of stem cells at the expense of differentiation. Conversely, induced increase in oxygen levels leads to premature neurogenesis (Lange et al., 2016).

We find a similar relationship between the hypoxia response and the differentiation state of different brain compartments in the *Drosophila* brain. A large part of the central brain comprises a fully differentiated nervous system, which controls larval behaviour. In the central brain there are only about one hundred neural stem cells or neuroblasts, which undergo self-renewing divisions to generate intermediate ganglion mother cells and secondary neurons for the adult nervous system. In contrast, in the optic lobes a pool of several hundreds of neuroepithelial cells arises through symmetric proliferative divisions. Neuroepithelial cells are then progressively transformed into optic lobe medulla neuroblasts at the end of larval development (Egger et al., 2007; Yasugi et al., 2010), which generate GMCs and neurons for the adult visual system. These neurons remain in a state of arrested differentiation until the second half of pupal life, when a wave of massive synaptogenesis starts (Melnattur and Lee, 2011; Chen et al., 2014). This indicates the existence of a mechanism that arrests differentiation of neural progeny during larval life. Hypoxia could be part of such a mechanisms and future experiments should reveal whether ambient hyperoxia, or local hyperoxia caused by ectopic tracheolation of the optic lobe result in altered mitotic activity and premature synaptogenesis.

In correlation with the observed differences in tracheolation, we found that cells in the optic lobe show a stronger hypoxic response as compared to cells in the central brain. The hypoxia biosensor provides a remarkable spatial resolution and it is possible to detect a differential hypoxia response in cells that are in close proximity to each other. We combined the ratiometric analysis with cell type specific markers and found that central brain and optic lobe neuroblasts appear to be the most hypoxic cell types in the developing larval brain. It suggests as documented also by other studies that lower oxygen is a condition that favours more undifferentiated and multipotent cell types (Eliasson et al., 2010; Lange et al., 2016). More committed neuronal progenitors such as ganglion mother cells and more differentiated cell types such as neurons and glial cells appeared to have a weaker hypoxia response. This suggests that a less hypoxic environment favours a more advanced state of cell differentiation. This is not the first time that such a model was suggested. Cipolleschi and colleagues proposed that the distance between a cell and a blood vessel in the stem cell hematopoietic niche could be an indicator of cell potency (Cipolleschi et al., 1993). A cell exposed to low levels of oxygen has higher multipotency and less fate commitment than a cell exposed to high oxygen. A similar relationship could be in place in the *Drosophila* larval optic lobe, where mitotic progenitor cells will be protected from oxidative damage by being segregated within a more hypoxic microenvironment, which becomes gradually more oxygenated as they advance in the differentiation process. In *Drosophila* a previous study by Homem and colleagues suggest that larval self-renewing neuroblasts initially use glycolytic pathways and only switch to oxidative phosphorylation at later pupal stages to initiate neuroblast shrinking and differentiation (Homem et al., 2014). Hence it would be interesting to investigate the relationship between neuroblast differentiation and tracheolation at pupal stages.

We provide evidence that oxygen levels as measured by the hypoxia biosensor decrease exponentially with the increase of the distance between a cell and its closest tracheole. While the great majority of cell types investigated here follow this model, we found some striking exceptions and most prominently the neuroepithelial cells of the inner proliferation centre. These cells are in close proximity of the densely tracheolated central brain but reveal an unexpectedly strong hypoxia response. Meanwhile, GMCs in the central brain show a much lower hypoxia response as compared to the predicted values. This prompts the question to what degree cell type and cell intrinsic mechanisms are responsible for a differential hypoxia response. Our study leads to the interpretation that the oxygen-dependent degradation of HIF-1α within a given cell depends mostly on its distance to tracheoles, but to a certain degree also on cell-type specific features. Among several possible explanations, we regard as probable the existence of cell-type specific differences in transcriptomes, metabolisms and the general capacity of given cell types to degrade the GFP-ODD of the biosensor.

We found that while neuroblasts adapt their hypoxia response to elevated ambient oxygen levels (hyperoxia), optic lobe neuroepithelial cells seem to have a more limited capacity to respond. This might be due to differential gene expression of hypoxia pathway genes in these two cell types. Indeed, by re-analysing cell type specific Polymerase II DamID data (Southall et al., 2013) we found that the transcriptome of neuroblasts, but not that of neuroepithelial cells, is highly enriched in hypoxia response genes.

Normally hypoxia triggers an adaptive response, which among other things upregulates glycolysis and angiogenesis (in mammals) or tracheolation (in insects). In *Drosophila*, this can be driven through different pathways, of which at least one is HIF-1α dependent (Li et al., 2013). In *Drosophila* larvae, hypoxia induces an increase in the length and branching of tracheoles in a matter of hours (Jarecki et al., 1999; Centanin et al., 2008). Here we reported that this does not happen in the optic lobe even after several days of larval life. On the contrary, our measurements indicate that the proportion of optic lobe deprived of tracheoles increases with age, suggesting the existence of a mechanism that mediates adaptation through enhanced glycolysis but without compensatory growth of tracheoles. The finding of the neuroepithelial gene catalogue not being enriched in genes of the cellular response to hypoxia appears to support this possibility and is, in turn, consistent with our hypothesis that the lack of tracheolation in the optic lobe is an adaptation to keep neural stem cells in a hypoxic compartment and as such an important feature of normal brain development. Hypoxia in the larval optic lobe will, in turn, promote proliferation and inhibit terminal differentiation of neurons.

## MATERIALS AND METHODS

### Fly stocks

Flies were raised on cornmeal medium at 25°C under light-dark cycles of 12:12 hrs light:darkness. Oregon R was used as wild type. The oxygen biosensor line used in this study contains the transgenes *ubi-GFP-ODD* (Misra et al., 2017) and *ubi-mRFPnls (BL-34500)*, both inserted on the second chromosome.

### Larval staging, dissections and immunostainings

Eggs were collected during a two hours period on apple juice agar plates. After 24 hrs freshly hatched larvae were transferred to standard cornmeal medium and placed in an incubator at 25°C with a 12:12 hrs light:dark cycle. Larval brains were dissected in 4% paraformaldehyde in 0.1M phosphate-buffered saline (PBS, pH 7.4), 0.5 mM EGTA, 5 mM MgCl and fixed for 18 minutes (including the time of dissection), washed with PBS containing 0.1% Triton X-100 3 times for 5 min and 4 times for 15 min. For immunofluorescent stainings, whole larval brains were incubated in primary antibodies overnight at 4°C. Incubations with secondary antibodies were also performed overnight at 4°C. Samples were washed at room temperature and mounted in Vectashield H-100 (Vector Laboratories). The following primary antibodies were used for specific protein labelling: mouse anti-Bruchpilot 1:50 (Nc82, DSHB; (Hofbauer et al., 2009)); mouse monoclonal anti-Discs large 1:30 (4F3, DSHB); guinea pig polyclonal anti-Deadpan 1:1000 (gift from A. Brand, Cambridge UK), mouse monoclonal anti-Prospero 1:10 (MR1A, DSHB; (Campbell et al., 1994)), rat polyclonal anti-Elav 1:20 (7E8A10, DSHB; (O’Neill et al., 1994)), and mouse monoclonal anti-Repo 1:10 (MAb 8D12, DSHB deposited by Goodman, (Alfonso and Jones, 2002)). The chitinous cuticle lining of the tracheal lumen was visualized either by its own autofluorescence under UV illumination or after staining with the fluorescent chitin-marker Calcofluor (1:200; Sigma). Appropriate secondary antibodies conjugated to Alexa fluorochromes were used: Alexa 405, Alexa 488, Alexa 568 and Alexa 633 all 1:200 (Molecular Probes).

### Hyperoxia and hypoxia treatments

Embryos were collected for two hours at 25°C on apple juice-agar plates and transferred to plates with standard food medium. For hyperoxia treatment, at 24 hrs after larval hatching (ALH) larvae were placed in a Modular Incubator Chamber (MIC-101; Billups-Rothenberg, Inc.), which was flushed with pre-mixtures of 60% oxygen in nitrogen (PanGas AG). At 84 hrs ALH, towards the end of the wandering stage, the larvae were dissected as described above. Oxygen concentration inside the chamber was monitored with an oxygen analyzer/monitor (Vascular Technology VTI-122 Disposable Polarography Oxygen Cell Catalog No. 100122). For the hypoxia treatment, larvae were collected at 60 hrs ALH and placed in the incubation chamber previously flushed with 5% oxygen pre-mixture in nitrogen for 24 hrs. Larvae under hypoxia tend to escape the medium, refrain from feeding and become delayed in development (Callier et al., 2013; Wong et al., 2014; Bailey et al., 2015). For this reason, we decided to use a shorter hypoxia treatment that was still long enough to observe a change in brain oxygenation with the biosensor.

### Light microscopy, image processing and analysis

For each brain, a single hemisphere was imaged using a 60x objective on a TCS Leica SPE-II laser scanning confocal microscope or with a 60x objective on an Olympus Fluoview FV300. Optical sections across the entire brain hemisphere were recorded at 0.6 µm intervals for tracheal surface measurement, at 1 µm intervals for the temporal ratiometric analysis with the hypoxia biosensor and at 2 µm for cell type specific ratiometric analysis with the hypoxia biosensor. The ratiometric anaylsis was performed with Fiji as described in (Misra et al., 2017). The macro and plugin can be found on Github: https://github.com/eggerbo.

Measurements of tracheolar length (Figure 2) were done separately in the optic lobe and in the central brain with Simple Neurite Tracer (Longair et al., 2011). TrakEM2 in Fiji (Schindelin et al., 2012) was used to segment the two volumes to be compared (optic lobe and central brain) following the neuroanatomical borders outlined by anti-Dlg staining. Reconstructions and quantifications were done in Fiji. Figures and illustrations were assembled using Adobe Photoshop 8.0, Adobe Illustrator 11.0 and Inkscape 0.92.

### 3D trachea map annotation and proximity analysis to brain cells

A pixel-resolution map of the tracheal system in the whole brain hemisphere was produced. To obtained this map, a stack of images containing the tracheal system were processed by background subtraction and smoothening. Then, a threshold was applied and the x,y,z coordinates of every pixel corresponding to every tracheole was stored to produce a digital map. Our ratiometric analysis provides ratiometric values for every nucleus n the brain hemisphere together with their x,y,z coordinates. Combining both data sets we calculated the Euclidean minimal distance from every nucleus in the brain hemisphere to the tracheal system. The analysis was performed in Python.

### Prediction of Ratio values based on tracheole proximity

The relationship between 1/Ratio values and proximity to tracheoles was best modelled by a decaying exponential function of the form:

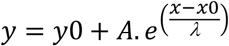

where λ is the length constant. λ, *y*0 and *A* are fitting parameters and *x*0 is a constant. We used the exponential function resulting from the best fit to the data to predict the value of hypoxia response according to tracheole distance. We used Fiji to represent these values for every nucleus in the brain with a colour scale (Figure 4 shows a single section of the resulting stack image map). Fits were preformed and analyzed with Igor Pro (Wavemetrics).

### Cell type specific analysis

To analyse the hypoxia biosensor in a cell type-specific manner ratiometric values were measured only for a specific cell type at a time. For this, a segmentation mask was generated for each major cell type in the brain using the fluorescent intensity signal corresponding to each cell type: Glia were identified by expression of Repo (mouse monoclonal anti-Repo); neuroblasts by the expression of Deadpan (guinea pig polyclonal anti-Dpn), ganglion mother cells by the expression of Pros (mouse monoclonal anti-Pros); and neurons by the expression of Elav (rat polyclonal anti-Elav). The inner proliferation center (IPC) and outer proliferation center (OPC) were segmented using the TrakEM2 software in Fiji following neuroanatomical borders outlined by anti-Dlg staining.

### Transmission electron microscopy

Brain samples for transmission electron microscopy were prepared from five wild-type (Oregon R) 96 hrs ALH larvae, according to the protocol detailed in (Talamillo et al., 2008). The brain was oriented and trimmed to obtain slightly tilted frontal views of one hemisphere. Ultrathin (50-60nm) sections were observed with a JEOL JEM 1010 electron microscope operated at 80kV. Several grids of each brain, each containing several sections, were observed. Images were taken with a Hamamatsu C4742-95 camera and processed with AMT Advantage and Adobe Photoshop.

### Statistical analysis

All data analyses and graphs were done using R/Bioconductor. Scripts for graphs can be found here: https://github.com/MartinBaccinoCalace. Biological replicates (n) are single brain lobes of different animals. For the optic lobe and central brain ratio values boxplot charts were created in such a way that each boxplot contained the normalized frequency values for seven replicates in a given bin. Student t-tests were performed when assumptions for parametric test were accomplished (normality using Shapiro-Wilk test and homoscedasticity using Levene’s test). If these assumptions were not achieved, nonparametric Mann-Whitney U tests were performed instead. Statistical significance was set at 0.05, 0.01 and 0.001. For tracheolation analysis Mann-Whitney U tests were performed and statistical significance was set at 0.05. Power analysis was conducted in RStudio to estimate minimum sample size for a power of 0.8 and α level of 0.05 for a two-sided student t test or Mann-Whitney U test.

### Gene enrichment analysis

A previous published dataset (Southall et al., 2013) was re-analysed to investigate hypotheses postulated in this paper. It contains cell-type specific gene expression catalogues of neural stem cells at third larval instar, obtained with an assay of RNA Polymerase II occupancy in either neuroepithelial cells or neuroblasts. The catalogues of genes expressed in either neuroepithelium or neuroblasts were curated to correct for gene annotation changes in the current version of the genome and the resulting catalogues were used to calculate whether they were enriched for the Gene Ontology terms in biological functions “cellular response to hypoxia” (GO:00771456) or “glycolytic process” (GO:0006096). To test for significance the hypergeometric test was used.

## ACKNOWLEDGEMENTS

We thank A. Brand, the Developmental Studies Hybridoma Bank and the Bloomington Stock Center for reagents and fly lines. We thank F. Meyenhofer for support on the image analysis pipeline and H. Akarsu for assistance with the R scripts. Special thanks go to S. Luschnig for the biosensor line and for valuable comments on the manuscript. We also thank the Sprecher laboratory for insightful discussions. M.B-C. was supported by Agencia Nacional de Investigación e Innovación, grant FCE_3_2013_1_100732. R.C. and M.B-C. were supported by Uruguay’s Programa de Desarrollo de las Ciencias Básicas. R.C. and DP received support from Uruguay’s Sistema Nacional de Investigadores. B.E. was supported by the Swiss University Conference (P-01 BIO BEFRI).

## AUTHOR CONTRIBUTIONS

M.B-C. Writing original draft, Investigation, Formal analysis, Visualization.

D.P. Writing-review and editing, Investigation, Formal analysis, Visualization.

R.C. Conceptualization, Writing-review and editing, Investigation, Formal analysis, Supervision, Funding Acquisition.

B.E. Conceptualization, Writing-review and editing, Investigation, Data curation, Supervision, Project Administration, Funding Acquisition.

